# A microbial community growth model for dynamic phenotype predictions

**DOI:** 10.1101/2022.12.15.520667

**Authors:** Andrew P. Freiburger, Jeffrey A. Dewey, Fatima Foflonker, Gyorgy Babnigg, Dionysios A. Antonopoulos, Christopher Henry

## Abstract

Microbial communities are increasingly recognized as key drivers in animal health, agricultural productivity, industrial operations, and ecological systems. The abundance of chemical interactions in these complex communities, however, can complicate or evade experimental studies, which hinders basic understanding and limits efforts to rationally design communities for applications in the aforementioned fields. Numerous computational approaches have been proposed to deduce these metabolic interactions – notably including flux balance analysis (FBA) and systems of ordinary differential equations (ODEs) – yet, these methods either fail to capture the dynamic phenotype expression of community members or lack the abstractions required to fit or explain the diverse experimental omics data that can be acquired today.

We therefore developed a dynamic model (CommPhitting) that deduces phenotype abundances and growth kinetics for each community member, concurrent with metabolic concentrations, by coupling flux profiles for each phenotype with experimental growth and -omics data of the community. These data are captured as variables and coefficients within a mixed integer linear optimization problem (MILP) designed to represent the associated biological processes. This problem finds the globally optimized fit to all experimental data of a trial, thereby most accurately computing aspects of the community: (1) species and phenotype abundances over time; (2) a linearized growth kinetic constant for each phenotype; and (3) metabolite concentrations over time. We exemplify CommPhitting by applying it to study batch growth of an idealized two-member community of the model organisms (*Escherichia coli* and *Pseudomonas flourescens*) that exhibits cross-feeding in maltose media. Measurements of this community from our accompanying experimental studies – including total biomass, species biomass, and metabolite abundances over time – were parameterized into a CommPhitting simulation. The resultant kinetics constants and biomass proportions for each member phenotype would be difficult to ascertain experimentally, yet are important for understanding community responses to environmental perturbations and therefore engineering applications: e.g. for bioproduction. We believe that CommPhitting – which is generalized for a diversity of data types and formats, and is further available and amply documented as a Python API – will augment basic understanding of microbial communities and will accelerate the engineering of synthetic communities for diverse applications in medicine, agriculture, industry, and ecology.

## 2 Introduction

Microbial communities are ubiquitous on Earth [1], and serve fundamental roles as eukaryotic symbionts [2, 3, 4] and pathogens [5, 6], industrial assets [7] and antagonists [8], and ecological agents of biogeochemical cycling [9, 10, 11]. Microbial communities are therefore essential to understand in fields as diverse as medicine [12, 13] and climatology [14, 15, 16]. The assemblage of microbes into an community, while inviting intimate competition [17], offers a few advantages that make it biologically favorable in most conditions. Firstly, diversification in a) genetic potential, b) biochemical vulnerabilities [18], and c) metabolic machinery [19, 20, 21] allows community members to grow in otherwise inhospitable conditions, such as nutrient-poor or toxic environments, by relying on well-adapted members. Secondly, specialization among the members enables the community to achieve more efficient production of vital nutrients [22] and greater growth rates [23] than is possible by isolated, autonomous, members. These interactions that guide community assemblage are predicated upon a metabolic economy, where the competitive community environment diversifies members into ecological niches that balance nutritional satiation and metabolic efficiency by leveraging cross-feeding [24, 25]. Metabolic cross-feeding (syntrophy) is essential to understand and engineer communities for desirable properties [26], such as bioproduction that exceeds the capabilities of monocultures [7] or sustainable biotechnology by coupling phototrophs and heterotrophs [27], yet it is difficult to experimentally measure.

One impediment to measuring syntrophy is that some metabolites do not detectably accumulate in the media, which obviates all but a few experimental methods, such as fluxomics isotope labeling [28]. A second difficulty, which proves to be more formidable, is that the combinatorial quantity of cross-feeding exchanges creates an untenable multitude of experiments to resolve all possible interactions in natural communities. These challenges are further compounded by the numerous metabolic phenotypes of each member that dynamically respond, even in clonal monocultures [29], to environmental conditions. Phenotype variability is necessary to understand community dynamics, and particularly to engineer members for production or exclusion, but this analysis adds another experimental dimension to an already challenging and expensive series of measurements.

Computational biology offers a few tools to resolve the metabolic nuances of microbial communities. Flux Balance Analysis (FBA) [30], for example, is a prominent framework for simulating cellular metabolism and has been applied in numerous software tools that seek to resolve metabolic interactions within microbial communities: e.g. BacArena [31], *µ*bialSim [32], dOptCom [33], SMETANA [34], CASINO [2], and MICOM [35], to name a few. FBA is attractive as a simple method for simulating metabolism that can operate without or, optionally, with experimental -omics data types [36] such as metabolomics [37, 38], proteomics [39], transcriptomics [40, 41, 42], or multi-omics [43] data. The under-determined linear problem of FBA, however, often lessens the precision and consistency of predictions and FBA moreover cannot natively capture phenotype variability within a given species metabolic model. Machine-learning algorithms [44, 45], in contrast, offer more exploratory flexibility for phenotype abundances from experimental data, but often lack the mechanisms that cultivate basic understanding and design principles. Differential equation models offer precise mechanistic explanations for predictions but has the converse problem [46] of frequently being stiff and insufficiently unbiased to flexibly explore unparameterized spaces. Fitting models, by contrast, as a final popular tool of computational biology, can be flexible enough to acquiesce new information from experimental data [47, 48, 49] while maintaining sufficient mechanistic resolution for basic understanding. Fitting methods have been accordingly applied to deduce kinetics coefficients [50] or phenotype expression [40] from experimental -omics data, but these methods do not convey time-resolved phenotype abundances within context of other experimental dimensions, such as metabolic concentration, that cultivate comprehensive understanding.

We therefore developed a global fitting model (CommPhitting) that deduces time-resolved information from experimental data: such as a) biomass of each community member and member phenotype; b) total concentrations of all metabolites that are used by the phenotypes; c) growth kinetics for each member phenotype; and d) conversion factors from each experimental signal to biomass that may accelerate the interpretation of future experiments. CommPhitting is a multi-species dynamic model of the community member phenotypes, where the phenotype flux profiles are rigorously defined through a sequence of FBA simulations – 1) a minimal growth is constrained (to prevent singularities); 2) carbonaceous non-phenotype consumption is minimized (to isolate the phenotype); 3) consumption of the phenotype source is minimized (to maximize the biomass yield); 4) optionally, specified phenotype excretions are maximized (to mirror experimental phenotypes); and 5) total flux is minimized (to find the most parsimonious flux profile) – that extends previously defined methods [40]. CommPhitting is generalized for various data types into a linear problem [51] that solves simultaneously to ensure that a global optimum is determined [52], and is concisely coded to execute within a minute on a personal computer.

We exemplify CommPhitting with an idealized 2-member community of model organisms (*Escherichia coli* and *Pseudomonas flourescens*), who exhibit complementary carbon preferences [53] but have not been investigated as a coculture. This benchtop community [54] demonstrated pivotal cross-feeding with acetate excreta from *E. coli* that was able to cultivate growth of *P. flourescens* in a non-viable media of maltose (Figure 1). CommPhitting fit the experimental growth data remarkably well, and the predictions of phenotype abundances and metabolic concentrations were validated with transcriptomics data from the NCBI GEO database [55] and in-house metabolomics measurements of our community, respectively. We further simulated BIOLOG data from this community, which revealed growth kinetics of this community for each condition and may have broad utility for predicting community behaviors in a variety of conditions. We anticipate using the knowledge from these simulations to guide the engineering of *E. coli* for community bioproduction. CommPhitting is available as an open-source Python module in the ModelSEEDpy and is operable with the KBase [56] ecosystem. We believe that this light-weight yet robust fitting model will illuminate nuances of community interactions and cultivate rational design of microbial communities for myriad fundamental and industrial applications.

**Figure 1:**
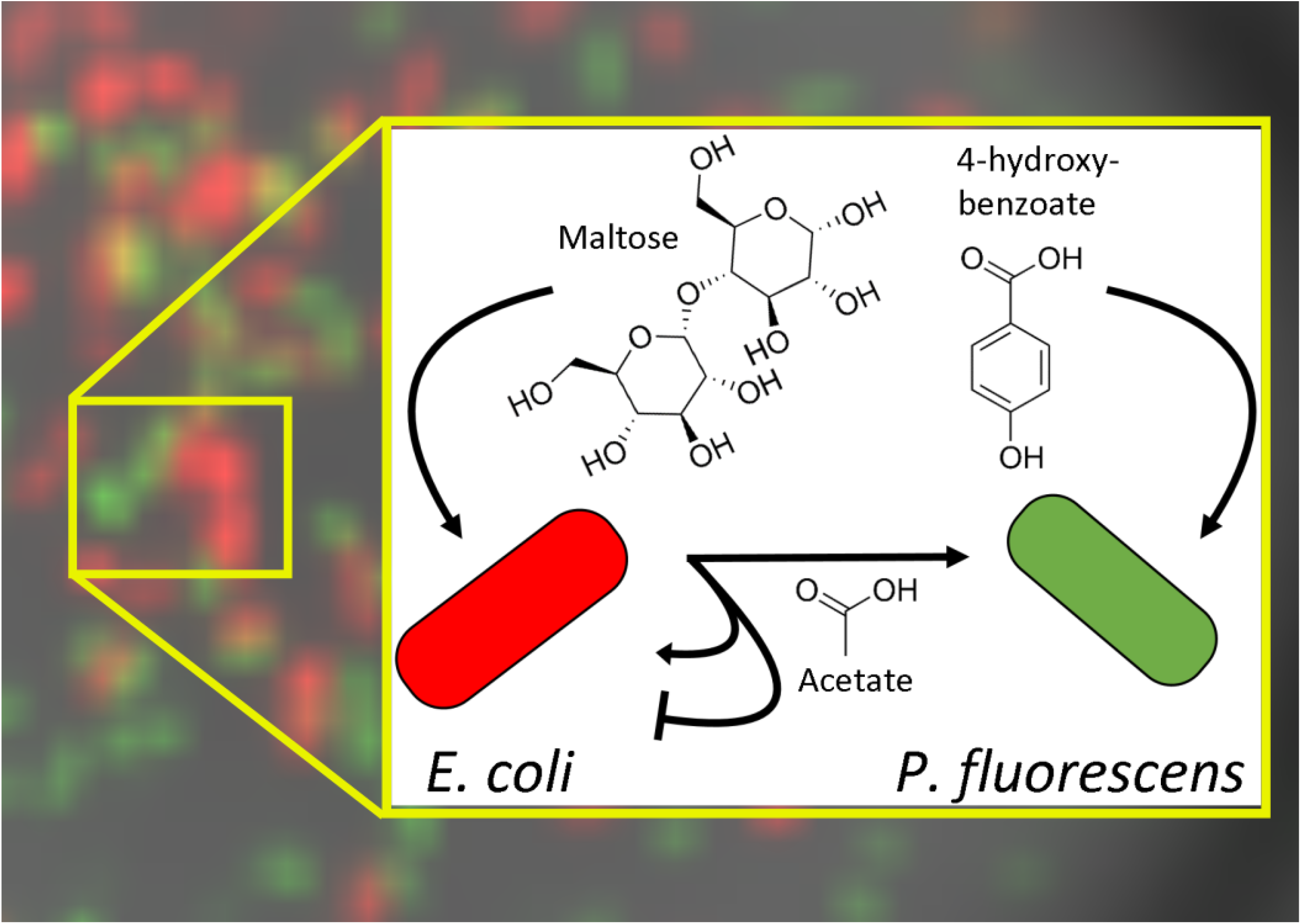
The primary metabolic exchanges that were experimentally elucidated and computationally modeled. The acetate byproduct of *E. coli* is the pivotal exchange of metabolic interest, where this source source subsists *P. fluorescens*’s existence in maltose and exhibits interesting dual effects upon *E. coli* as both a secondary carbon source and a growth inhibitor. Additional experiments revealed that *E. coli* is reluctant to grow in pure acetate and most often fails to grow at all, particularly when in cocultures with *P. fluorescens*.

## 3 Methods

The CommPhitting method transforms the traditional ordinary differential equations that define microbial growth and chemical activity into a mixed integer linear optimization problem (MILP). The MILP variables, constraints, and parameters are described in Table 1, and the MILP objective minimizes error in predicted growth, versus experimental growth, and minimizes overfitting. The processes and calculations that constitute the CommPhitting MILP are detailed in the following sections.

**Table 1:**
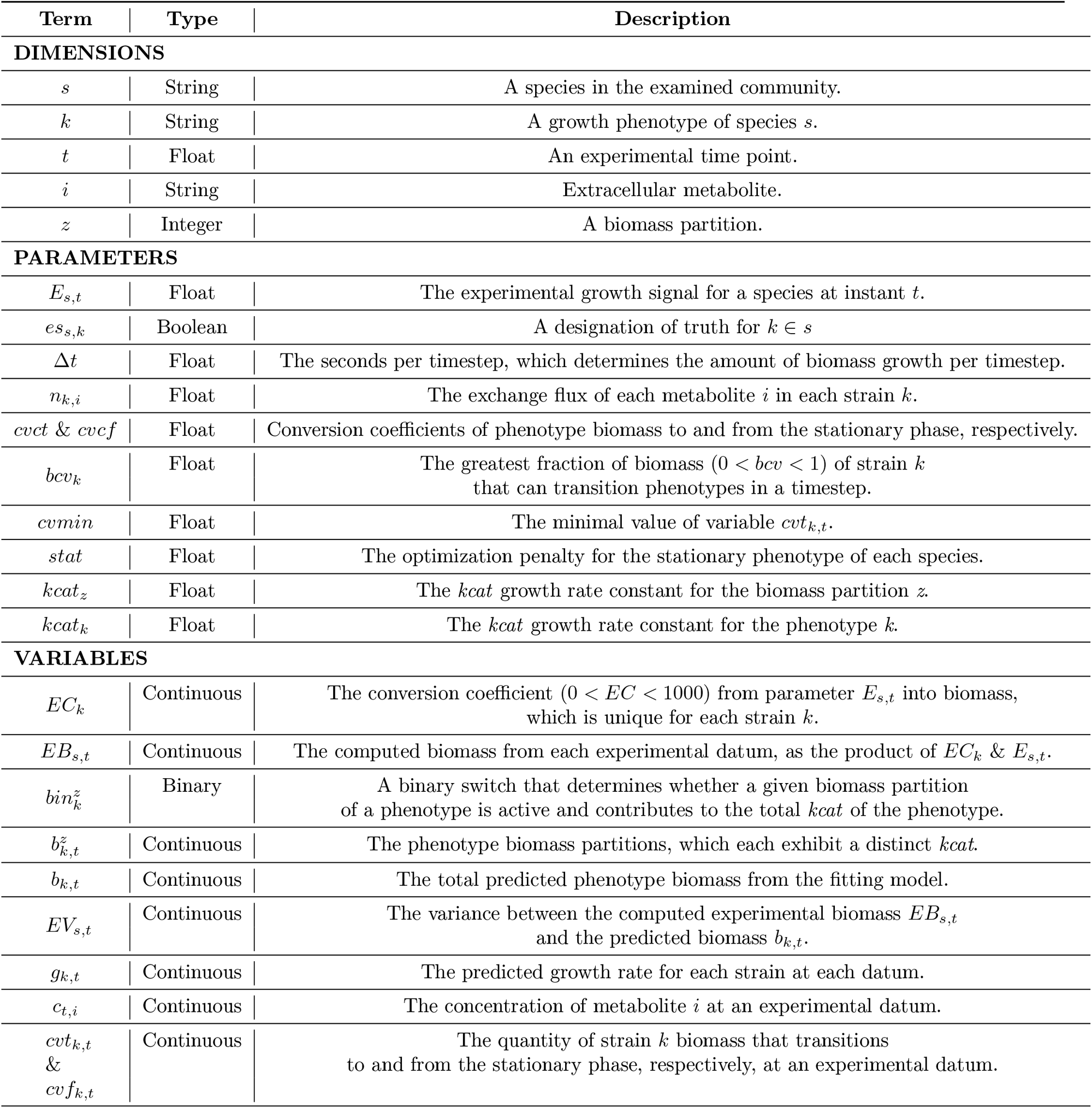
A glossary of dimensions, parameters, and variables that comprise the fitting model.

### 3.1 Metabolic phenotypes

The first step in constructing the CommPhitting MILP defines the metabolic phenotypes for each community member strain, withstanding the non-growing stationary phenotype. Experimental genomes of each member are reconstructed into genome-scale metabolic models (GEMs), where the ModelSEED pipeline [57] is advantageous among reconstruction tools for guaranteeing model consistency and compatibility (e.g. matching IDs for extracellular metabolites). The GEMs are then manipulated into specified phenotypes through the following sequence of FBA simulations.

1. The biological and environmental conditions of the GEM are constrained. A minimal biomass growth is defined; hydrogen consumption is prohibited, which forces consumption of the specific carbon source for growth; and oxygen consumption is limited to the total consumption of the defining phenotype carbon source(s), although, this threshold can fluctuate to accommodate the specific species biology and to properly regulate overflow metabolism.
2. The total influx of carbon compounds is minimized, excluding the defined phenotype carbon source(s) and any compounds with unknown formula. This optimally focuses on the metabolism that corresponds with consuming the designated carbon source(s) of the phenotype. The resultant fluxes from this minimization are fixed to the model and are unchanged in subsequent optimizations.
3. The total flux of the phenotype carbon sources is minimized, while fixing the biomass growth flux, which maximizes biomass yield from the designated carbon source(s). This presumably identifies the most efficient and biologically desirable nutrient consumption and byproduct excretion profile for each phenotype. The resultant carbon source fluxes from this minimization are fixed to the model.
4. Optionally, the excretion flux for assigned excreta of a phenotype (where experimental evidence is available) is maximized, and then the excreta flux is fixed to the model.
5. The final step applies parsimonious FBA (pFBA). This FBA method minimizes the sum of all fluxes at a fixed biomass growth and thereby finds the simplest flux solution to reach the prescribed growth. This pFBA simulation presumably emulates the efficiencies that evolution found for the simulated organism. The non-zero exchange fluxes from this final optimization becomes the CommPhitting representation of the specified phenotype.

The above five-step sequence is repeated for each specified phenotype of all species in the community. This robust method creates an irreducible metabolic profile for each phenotype that fosters pure phenotype predictions by CommPhitting.

### 3.2 Constraints

Each of the following linear constraints of CommPhitting captures a distinct aspect of community biology with the previously defined phenotypes.

#### 3.2.1 Biomass abundances

The experimental biomass (*EB*_*s,t*_), over all time points *t* ∈ *T*, is determined by converting the experimental fluorescence signal (*E*_*s,t*_) via a unique coefficient (*EC*_*s*_) for each member *s* ∈ *S*

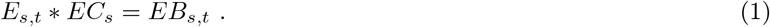

The final *E*_*s,t*_ is limited to be within 30% of the estimated biomass to consume 90% of the carbon source(s) according to the phenotype metabolic profile, which firmly anchors the *EC*_*s*_ conversion to data. The variance (*EV*_*s,t*_) between the converted experimental biomass and the predicted biomass *b*_*k,t*_ is determined for each phenotype *k* ∈ *K*

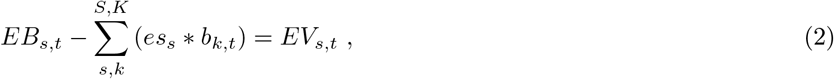

where only the phenotypes of each respective species are considered, per the binary *es*_*s*_ variable. The predicted biomass *b*_*k,t*_ is further partitioned into five sub-groups that are weighted along a gradient

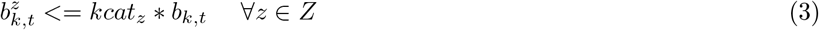

and that each exhibit a distinct *kcat*_*z*_ value. The expression of these biomass partitions is coordinated by the respective binary variable 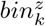 for each partition through the

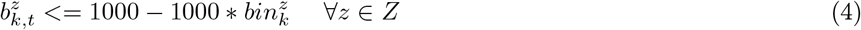

and

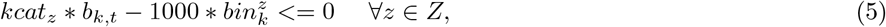

where one of the biomass partitions must be expressed

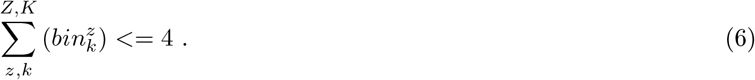

The *kcat*_*k*_ value of each phenotype is estimated by adjusting the *kcat*_*z*_ distributions and utilizing an exterior loop that changes the *kcat*_*z*_ values for each *z* ∈ Z partition (e.g. [100, 10, 1, 0.1, 0.01] in the first loop, [4, 2, 1, 0.5, 0.25] in the second loop, [1, 0.5, 0.25. 0.125, 0.0625] in the third loop),

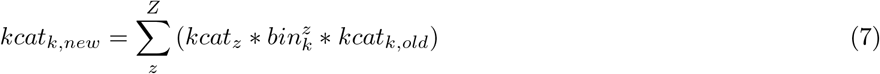

where *kcat*_*k,old*_ is guessed for the first loop and the solved *kcat*_*k,new*_ is substituted into the next loop as *kcat*_*k,old*_ for further refinement. This approach approximates *kcat*_*k*_ values for each phenotype while maintaining linearity of the growth constraint in eq. (9).

#### 3.2.2 Phenotype transitions

The biomass change over time is calculated for non-stationary (growing) phenotypes

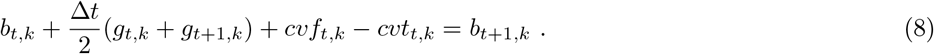

The constraint consists of terms for the current biomass (*b*_*k,t*_), the biomass growth rate *g*_*k,t*_ as the biomass derivative 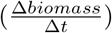, the biomass in the next timestep (*b*_*k,t*+1_), and the net transition of biomass to non-stationary metabolic phenotypes *cvf*_*k,t*_ − *cvt*_*k,t*_. The growth rate is constrained

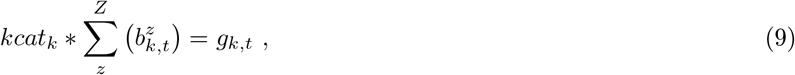

as the product of the current biomass abundance *b*_*k,t*_ and the growth rate constant *kcat*_*k*_, which reflects 1^*st*^-order kinetics with respect to biomass abundance and zero-order kinetics with respect to substrate concentration. Every species is defined with a stationary phenotype (not growing) that mediates and subtly delays phenotype transitions to reflect the inherent lag in pathway expression changes. Further, the stationary phenotypes allow each species to cease growth once the defined carbon source(s) is exhausted. Future biomass for the stationary phenotype is analogous to eq. (8),

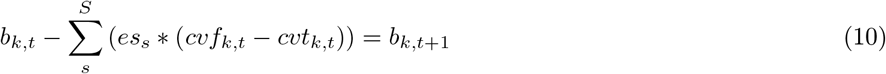

except that a) *g*_*k,t*_ = 0 by definition; b) the net transition to non-stationary phenotypes is negative; and c) the net transition to all of the many non-stationary phenotypes are summed. The fraction of biomass that can transition is constrained

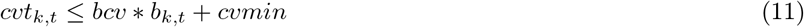

to greater than a minimum limit (*cvmin*) and lesser than a fraction of biomass (*bcv*) above this minimum.

This constraint is an application of Heun’s integration method [58, 59], which – for an arbitrary function *y*, its derivative *y*^*′*^, and a timestep ∆*t* – is defined as

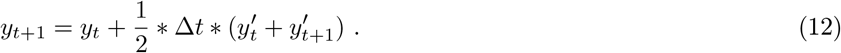

Heun’s method is a 2^*nd*^-order Runge-Kutta formulation [60] that captures dynamic changes with high numerical accuracy [61] while maintaining a linear formulation that is amenable with our MILP.

#### 3.2.3 Concentrations

Future concentrations of substrates (*c*_*t*+1,*i*_) for all *i* ∈ *I* substrates are constrained

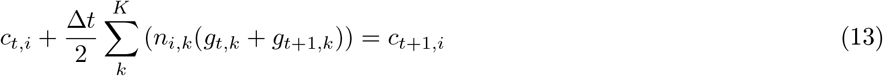

with Heun’s method from eq. (12). This implementation includes the current concentration (*c*_*t,i*_) and concentration changes that are the product of ∆*t* and the inner product of the metabolite flux (*n*_*i,k*_) from the phenotype flux profile (section 3.1) and the biomass growth rate for a member.

### 3.3 Objective

The objective expression

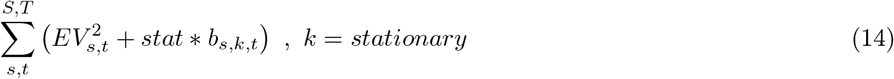

defines the sum of variance (*EV*) from eq. (2) – between the predicted (*b*) and experimental (*EB*) biomass – and the sum total amount of biomass in the stationary phenotypes. This objective is minimized to optimally fit and recapitulate the experimental growth, and minimally overfit from abusing stationary phenotypes or excessively transitioning phenotypes, where the stationary phenotype mediates phenotype transitions in eq. (8).

### 3.4 E. coli-Psuedomonas competitive community

#### 3.4.1 Experimental Methods

The exemplary *P. fluorescens* SBW25 and *E. coli* (minimally modified) MG1655 community was selected for several reasons. Firstly, these members are model organisms that encompass a wide range of disciplines, such as a) the human microbiome, b) bioproduction, c) synthetic biology, d) the rhizosphere, e) agriculture, and f) ecology. Secondly, despite these members being robustly studied individually, they have not been studied as a coculture, which offers new experimental information and mitigates biases and preconceptions in hypothesis development and results interpretation.

The coculture was created through the following protocol. The *P. fluorescens* and *E. coli* strains were purchased from ATC, were stored at − 80^*◦*^C before preparation for electrocompetancy, and were transformed with a plasmid to constitutively express either mNeongreen or mRuby2 fluorescent proteins (GFP and RFP), respectively [62]. Transformed cells were freshly streaked on a LB agar plate with appropriate antibiotics from − 80^*◦*^C glycerol stocks, and were incubated overnight at 30^*◦*^C. A single colony from the plate was picked, placed into liquid LB (Lennox) broth with the antibiotics, and shaken @ 250 RPM overnight at 30^*◦*^C. The 2 mL overnight culture was pelleted (4000x g for 10 min), the supernatant was removed, and the cells were resuspended in 1 mL of M9 media that contains no carbon source. This washing sequence was repeated twice. A 20 *µ*L aliquot of the washed cells was combined with 2 mL of M9 media that contained the appropriate carbon source for each strain – 10 mM D-maltose for *E. coli* and 6 mM 4-hydroxybenzoate for *P. fluorescens* – and was shaken @ 250 RPM for at least 16 hours. These overnight cultures were washed twice, following the same procedure as the overnight culture. These cells were finally analytically examined via optical density (OD 590 nm) and fluorescence using a plate reader (Hidex).

The aforementioned M9 cultured cells were mixed in fresh M9 media to achieve the desired initial cell ratio at OD 0.1 (590 nm) and carbon source concentration/ratios. A 200 *µ*L aliquot of the cell mixture was then added to wells of a sterile, black wall & clear bottom, 96-well imaging plate (Costar). The 96-well plate was then added to the Hidex plate reader and was prewarmed to 30^*◦*^C. The cells were shaken orbitally at 900 RPM in the plate reader for, at least, 24h while being measured for optical density (600 nm), red fluorescence (544 excitation, 590 emission), and green fluorescence (485 excitation, 535 emission) every 10 minutes.

We utilized two methods to accurately disentangle the composition of a liquid coculture: 12^*◦*^C and 37^*◦*^C temperature variability and fluorescence reporter with the green- and red-fluorescence proteins, for *P. fluorescens* and *E. coli* respectively. The precision and mutual validation of these methods is depicted through a combined liquid culture and agar plating experiment (Figure 2), which supports their application in our study as a resolving mechanism for the community members. In this qualitative experiment, 2 mL of M9 media containing either maltose, 4-hydroxybenzoate, or acetate are seeded with mono- or cocultures of *E. coli* and *P. fluorescens*to 0.1 OD. After 24h growth with shaking at 30^*◦*^C, the liquid cultures are diluted into M9 media and streaked out on LB agar plates. The plates are placed at three different temperatures to enable selective growth of specific organisms. Specifically, *P. fluorescens*grows perceptively after 60h at 12^*◦*^C but does not grow at 37^*◦*^C, E. coli. shows the opposite growth trend, and both organisms grow robustly overnight at 30^*◦*^C. The results and significance of this experiment are discussed below, but the temperature dependent plating method confirms that fluorescence changes observed in coculture accurately represent changes in each organism’s growth/abundance.

**Figure 2:**
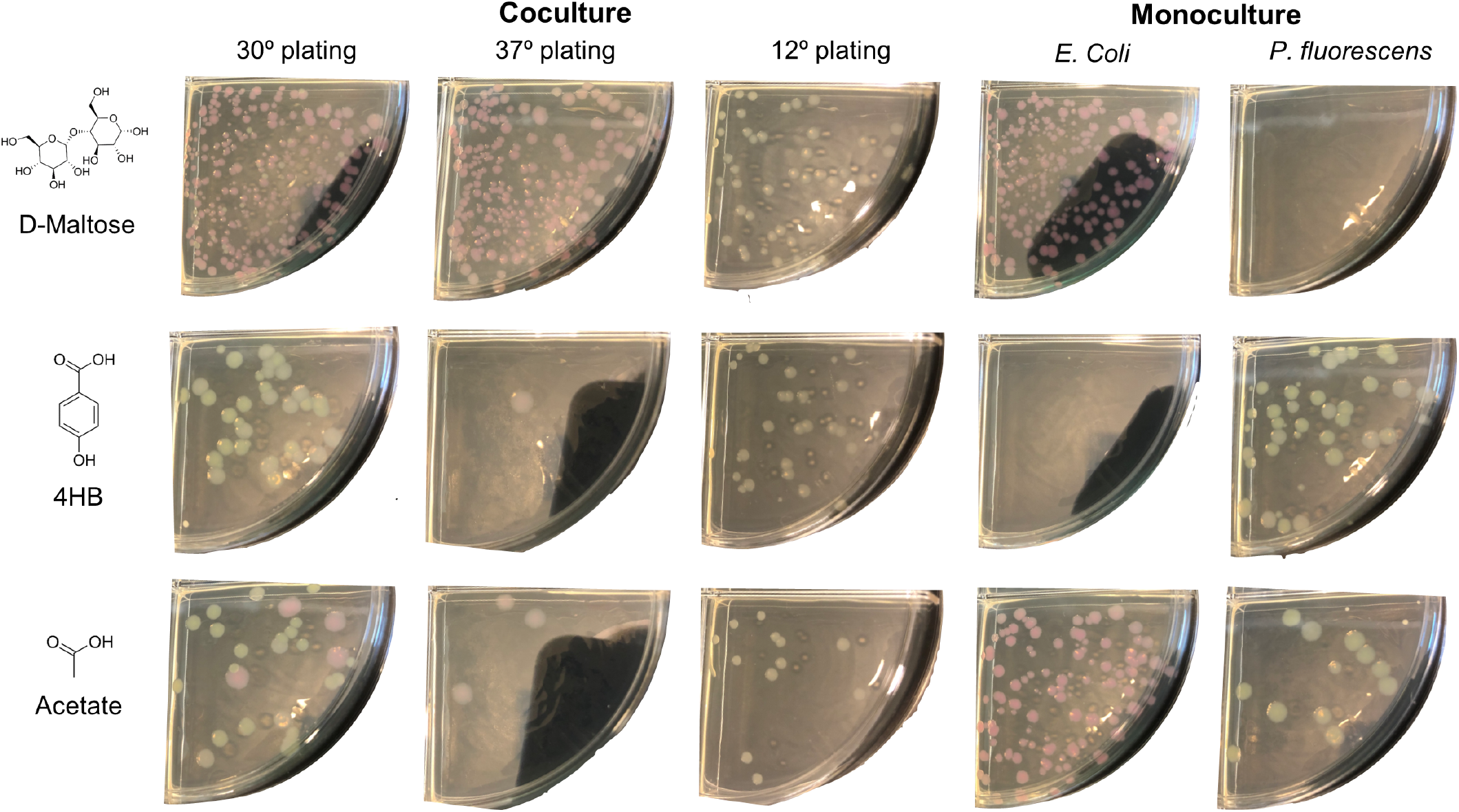
Organism abundance after co- and monoculture experiments in M9 media containing different carbon sources is visualized via plating at select temperatures. Coculture growth patterns replicate those observed in 96-well plate reader assays, confirming that fluorescence reporters accurately represent each organism’s growth/abundance.

#### 3.4.2 Adapting the formulation for growth data

The general formulation in Sections 3.1-3.3 was tailored to this particular 2-member community. First, the signal to biomass conversion constraint in eq. (1) was adapted to this community system

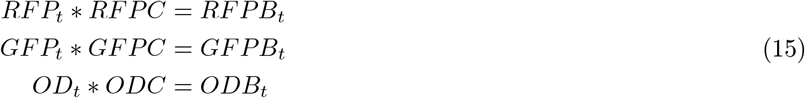

where the *RFPC, GFPC*, and *ODC* conversion factors were defined for each member and each *RFP*_*t*_, *GFP*_*t*_, and *OD*_*t*_ experimental signal. Second, the variance constraint in eq. (2) is adapted for each experimental signal

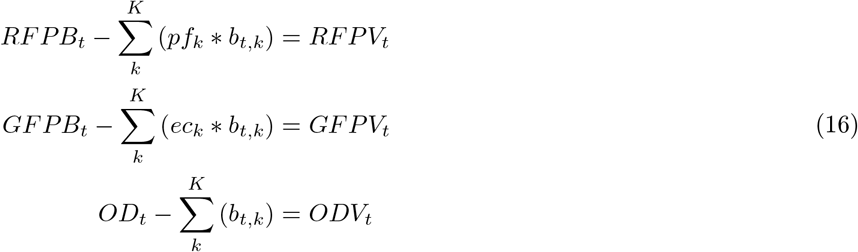

where *pf*_*k*_ and *ec*_*k*_ are binary variables that filter for *Psuedomonas* and *E. coli*, respectively. Third, the objective function of eq. (14) is defined for these members

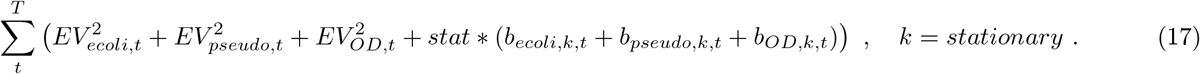

Finally, the data is filtered to remove timepoints beyond when the OD signal plateaus. This filtering mitigates the effects of fluorescent proteins over-production once cells enter their stationary phase and stop growing, by focusing on the lag and growth phases. These data could also be filtered by phenotype-specific conversions of the flourescence signals into biomass abundance, instead of species-specific conversion. Anomalous timesteps or datums can also be removed ad hoc.

The *P. fluorescens* and *E. coli* genome-scale metabolic models were constructed in KBase Narrative 93465 by leveraging RAST to annotate the genomes [63] and then applying the ModelSEED pipeline [57]. The *E. coli* model derived from the ASM584v2 experimental genome assembly and was gapfilled with acetate and maltose carbon sources. The *P. fluorescens* model derived from the ASM161270v1 experimental genome assembly and was gapfilled with acetate and 4-hydroxybenzoate carbon sources. We attempted to use published metabolic models for *E. coli* [64] *and P. fluorescens* [65], however, these models were a) not sufficiently responsive to the phenotyping methods of 3.1 and b) were developed by different groups, which introduced incompatibilities when simulated together in a community.

## 4 Results and Discussion

### 4.1 New data fitting method for iterative experimental design

The CommPhitting method, as detailed in Section 3, was developed to resolve phenotype abundances in microbial communities while accommodating diverse experimental data. The logical workflow of the model is illustrated in Figure 3. First, the experimental data of a trial is parsed and parameterized into a dynamic optimization model of the predicted metabolic growth phenotypes for all community members. Second, this global model is simulated to discover the variable and parameter values that best recapitulate the experimental data while minimizing overfitting. Finally, the simulation predictions – namely phenotype abundances, metabolite concentrations, and growth kinetic parameters – are expressed and exported as figures or spreadsheets.

**Figure 3:**
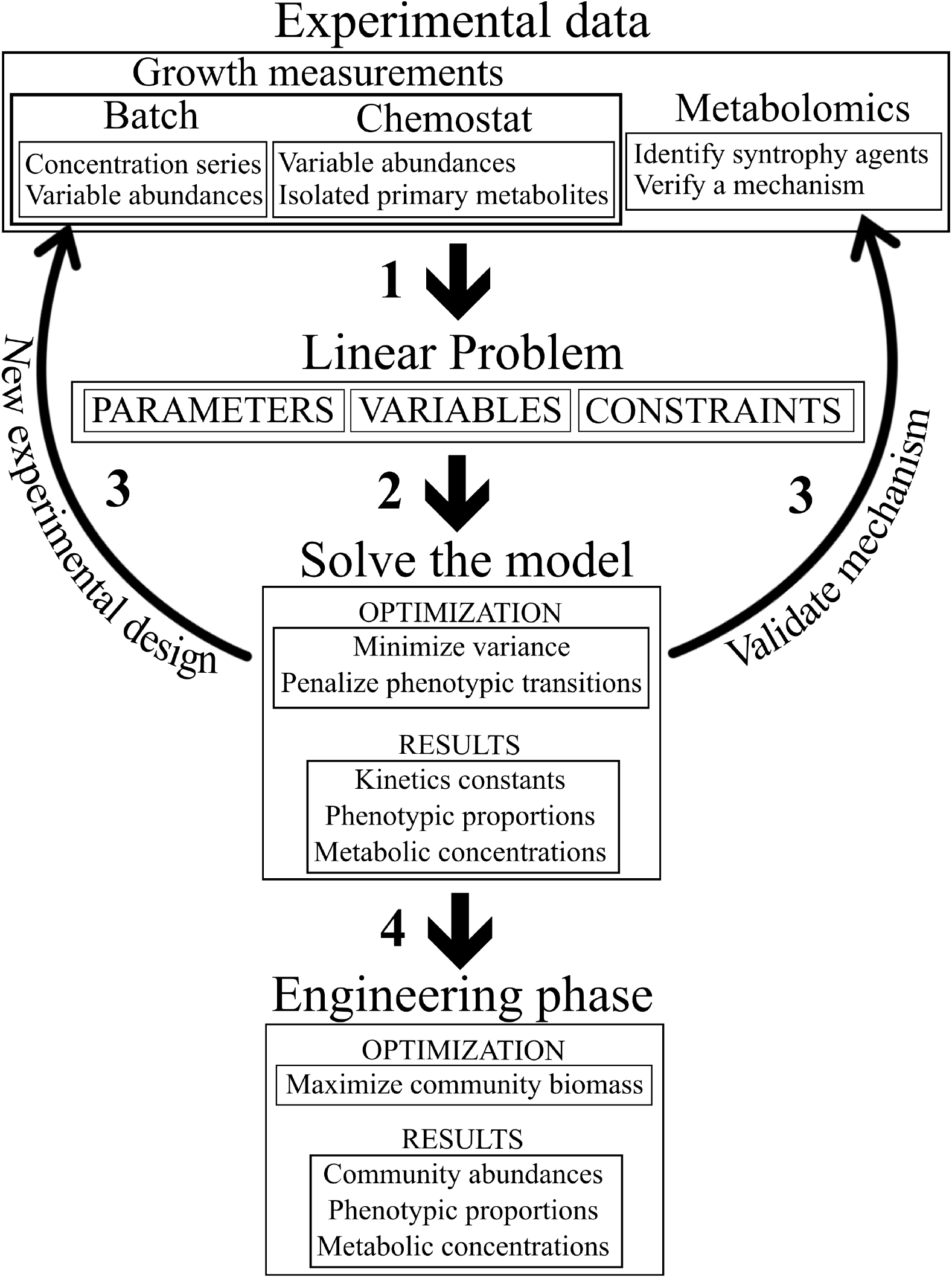
A workflow of the fitting model. **Step 1**: experimental data – from growth and possibly metabolomics measurements – is parsed into a MILP problem that consists of the parameters and variables that are detailed in Table 1 and the constraints that are explained in Section 3. **Step 2**: the linear problem is executed with the objective function of eq. (14). **Step 3**: the simulation results are interpreted to either identify experimental changes that will improve modeling fit or to propose select additional omics measurements that can further solidify mechanistic insights from the fit. Steps 1-3 repeat until a satisfactory fit and mechanistic resolution is achieved. **Step 4**: the fitted model can be used in a forward design-phase, instead of a purely retrospective fitting-phase, by replacing the objective function of the fitted model with one that maximizes community growth or targets other potential ecological objective functions. The system of steps 1-4 create an integrated method for gleaning mechanistic insights of microbial communities and then immediately using these insights to rationally design a community with desirable activity.

CommPhitting is designed as an open-source Python package – which leverages the Optlang optimization module [66], the GLPK solver, and ReadTheDocs documentation – to make the method as broadly accessible as is possible. The CommPhitting package further offers robust graphing capabilities through a simulation parameter that can consolidate the high-dimensional simulation results without requiring user proficiency with data science methods, which accelerates discovery by immediately gleaning biological insights after each simulation.

### 4.2 Simulation insights

The CommPhitting simulations of our 2-member community fit the growth experimental data in Figure 4 remarkably well, and thus instills confidence in the model predictions. A slight anomaly in the experimental data of the maltose + 4HB trials is compensated by CommPhitting in Figure 5, which demonstrates is flexibility while also communicating an area for experimental refinement. The simulations in maltose 6 elucidated the pure expression of the acetate phenotype in *Pseudomonas*, which kept the acetate concentration below 2*mM* throughout the simulation per Figure 7, while dynamic expression of *E. coli* between its three defined phenotypes: maltose, acetate, and stationary. The consumption of acetate is primarily predicted by *Pseudomonas*, but *E. coli* briefly becomes the greater acetate consumer as it begins to exhaust maltose and reaches the stationary phase. The contrast between *E. coli* in the coculture and as monoculture in Figure 6 exhibits a few interesting differences that illuminate the effect of *Pseudomonas* on *E. coli*. The first observation is that *E. coli* spends much more time in its stationary phenotype throughout the time-course, perhaps because of the inhibitory effects of acetate upon its growth, and *E. coli* enters the stationary phase a few hours sooner than in a coculture. A second observation is that *E. coli* reluctantly delays acetate consumption until the very end of the simulation, although it ultimately consumes more. A third distinction between *E. coli* as a monocultural versus as a coculture with *Pseudomonas* is the predicted *kcat*_*acetate*_, where the monocultural value is 8.89 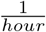 while the cocultural value is 0.23 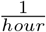. The lesser growth rate, or greater consumption of acetate without growth, may be a competitive adaptation of *E. coli* in the coculture to starve a competitors by consuming its food without necessarily investing the resources to grow itself.

**Figure 4:**
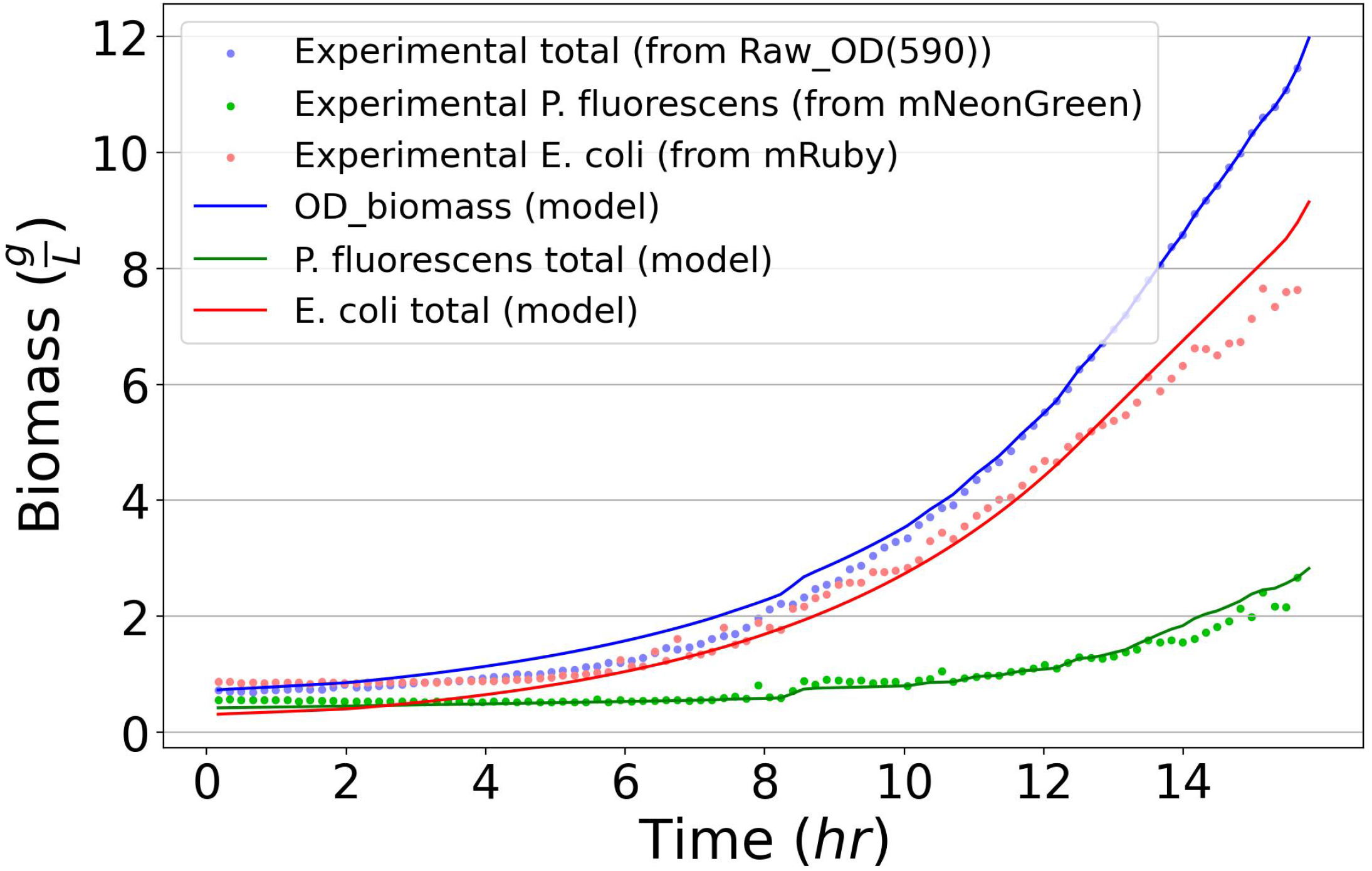
The converted experimental and predicted biomasses of the total community (OD), *E. coli* (RFP), and *P. flourescens* (GFP) for the coculture experiment on maltose media (Table S1). Tight agreement between the experimental and predicted biomass values improves confidence in the predicted community behaviors and underlying chemical parameters.

**Figure 5:**
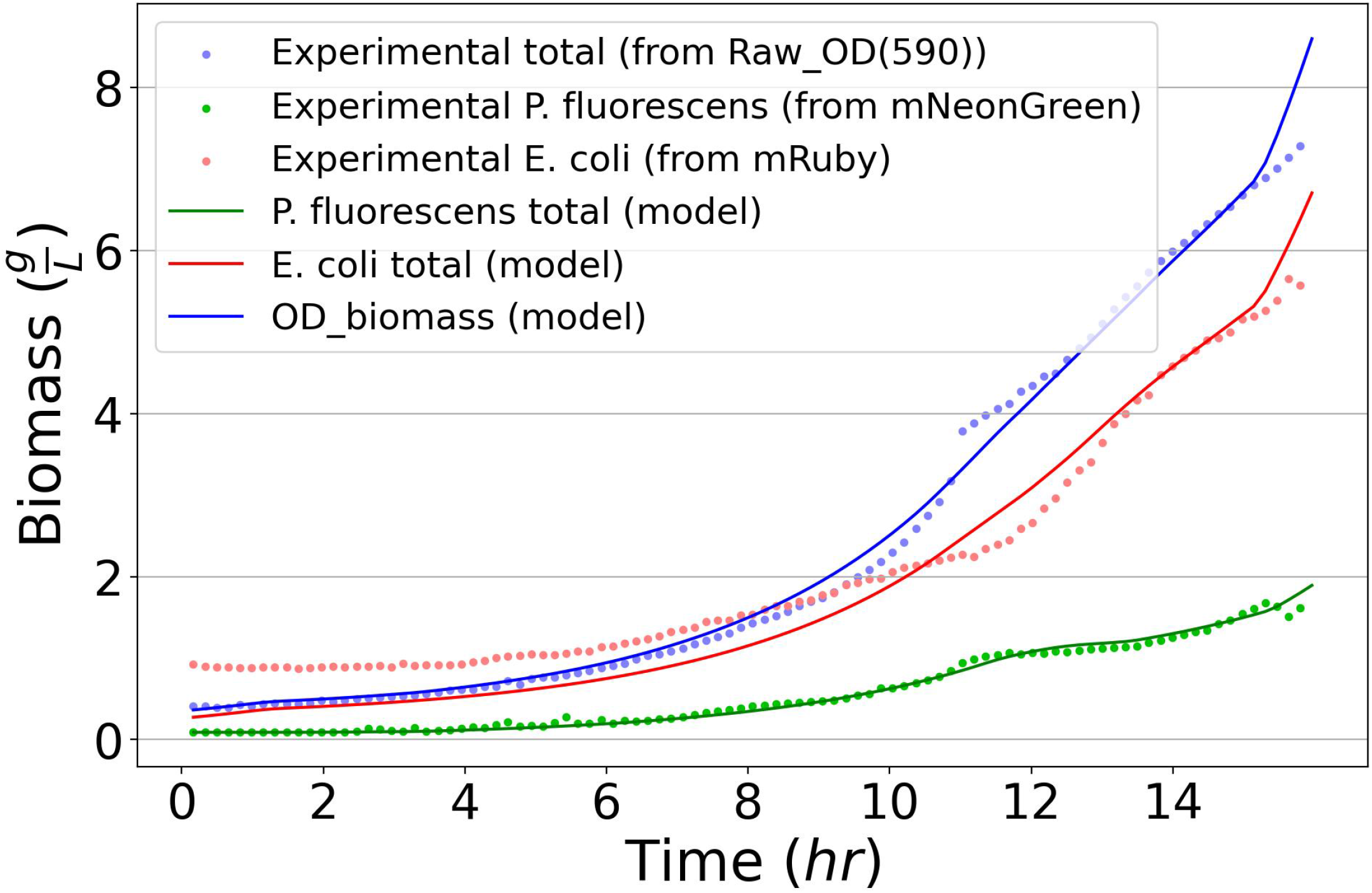
The converted experimental and predicted biomasses of the total community (OD), *E. coli* (RFP), and *P. flourescens* (GFP) for the coculture experiment on maltose media (Table S1). Tight agreement between the experimental and predicted biomass values improves confidence in the predicted community behaviors and underlying chemical parameters.

**Figure 6:**
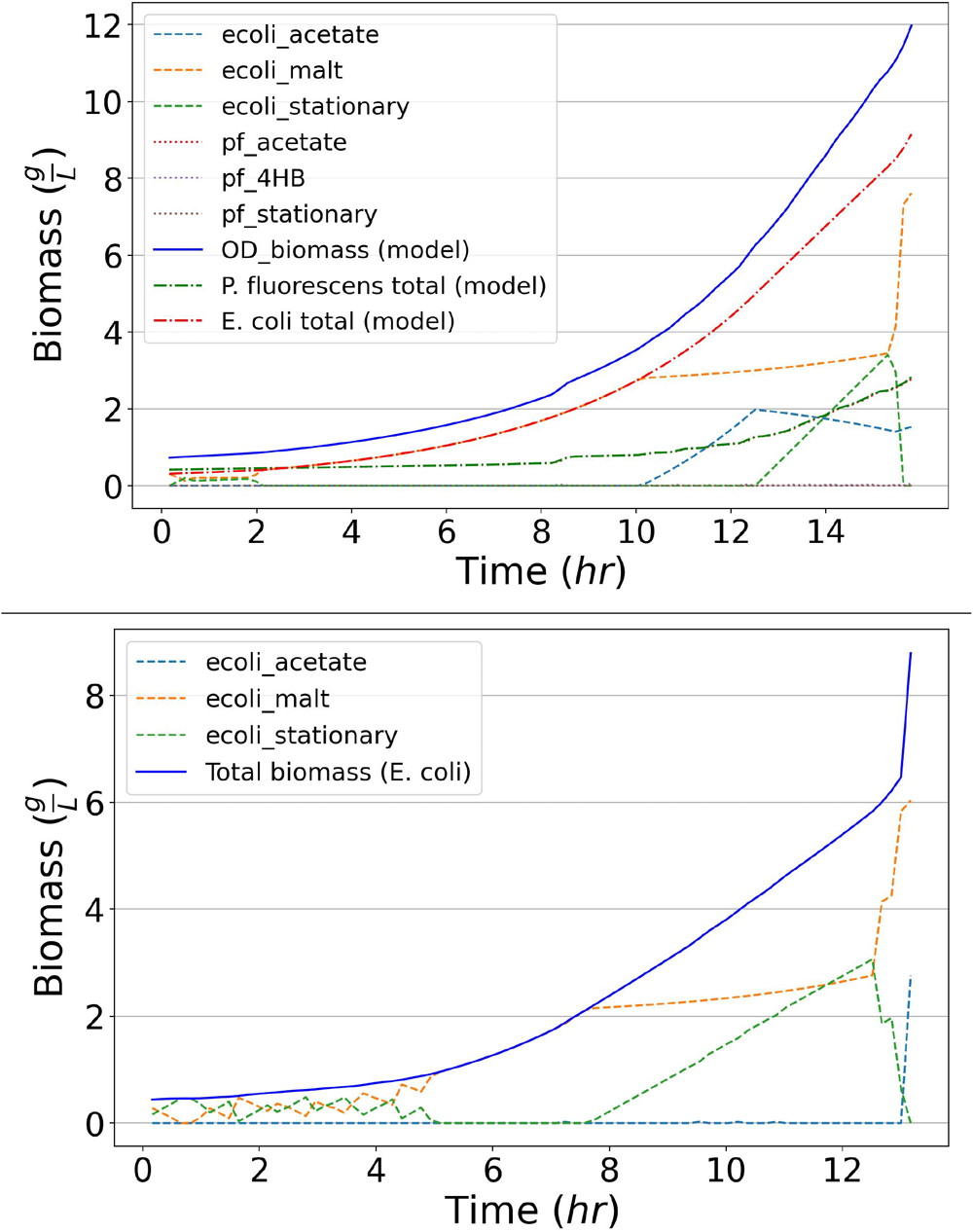
The phenotype abundances for an experiment with 5*mM* of maltose as the sole carbon sources. The top figure depicts the 1:1 coculture, where *E. coli* outcompeted *P. fluorescens*; however, *P. fluorescens* managed to subsist on the acetate excreta from *E. coli*. The bottom figure depicts a *E. coli* monoculture in maltose, where it reaches the same final biomass but consumes 50% more acetate, albeit very reluctantly. The monoculture interestingly also exhibited much more stationary phenotype during the lag phase than *E. coli* in the same conditions as a coculture. A *P. fluorescens* monoculture figure is not depicted since *P. fluorescens* exhibited no growth.

**Figure 7:**
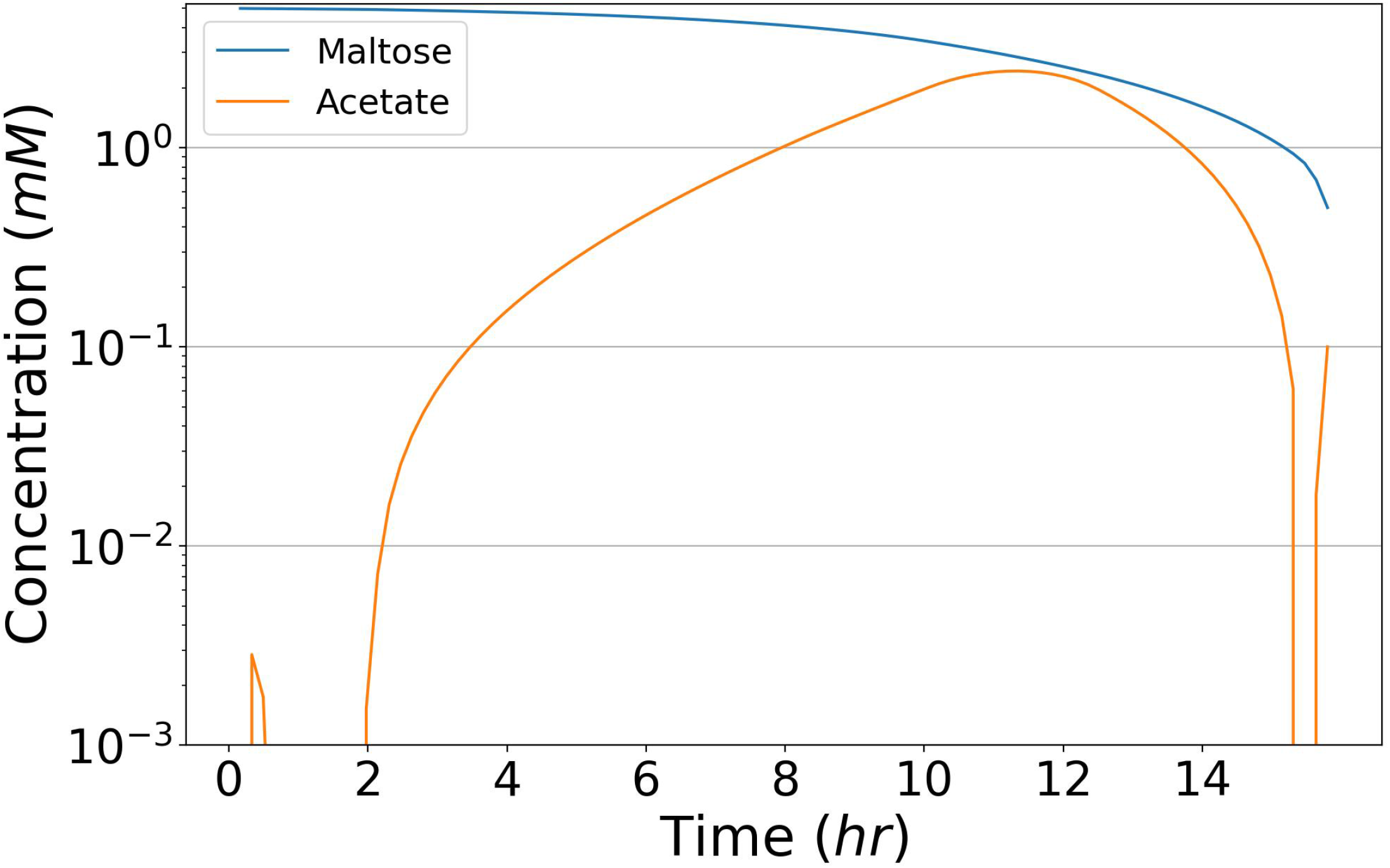
The concentrations of primary carbon sources and the predominate cross-feeding agent acetate from the coculture trial of Figure 6. The dynamic production and subsequent consumption of acetate elicits the unique information that can be derived from CommPhitting simulations. The peak acetate concentration at 2.4*mM* corroborates with the peak magnitude from the metabolomics data at the same OD value, which validates the concentration mechanisms of CommPhitting.

### 4.3 Validation

The CommPhitting simulations predicted minimal *E. coli* acetate consumption during coculture with *P. fluorescens*. This motivated our efforts to find *E. coli* mutants that lack this acetate absorption phenotype in order to corroborate: a) our experimental evidence that acetate is the predominate cross-feeding agent; and b) the prediction that the *E. coli* acetate phenotype is minimally expressed. We purchased mutant knock-outs from the *E. coli* Genetic Stock Center [67] for the four proteins – PoxB, ACS, Pta, and AckA – that contribute directly to acetate metabolism in MG1655. Assessing mutant growth in an acetate mono- or co-culture revealed that only the ∆Pta mutant terminated all acetate consumption; however, because this mutant excretes lactate instead of acetate [68] during sugar metabolism, *Pseudomonas* growth was accentuated by the coculture with this mutant since lactate is a more energetically favorable substrate than acetate. This study affirms that acetate is the primary mediating compound, where *Pseudomonas* responded profoundly to the one mutant whose acetate metabolism was disrupted, and inspires additional simulations of this community with new potential metabolic growth phenotypes, such as lactate, that explore other potential syntrophic interactions.

The peak acetate concentration in Figure 7 is furthermore validated with metabolomics data. The peak concentration in our simulation of 2.4*mM* corroborates with the peak concentration from the metabolomics data of 1.6*mM* at the same OD of 1.1. The experimental concentration at this OD is extrapolated between the enclosing metabolomics points at OD’s of 0.34 and 1.64; hence, the interpolated concentration embodies some error that may encompass the 50% excess of the predicted value, and it is possible that the predicted and experimental concentrations exhibit an even better match.

## 5 Conclusion

The ubiquity and uniqueness of microbial communities offer untold potential for basic understanding and technological advancement in diverse fields. This potential, however, remains hindered by knowledge gaps of phenotype dynamics and the interspecies exchanges that emanate from the combinatorial complexity of community interactions, which becomes experimentally intractable. Computational efforts to illuminate these gaps often improperly balance mechanistic resolution with data flexibility, and therefore fail to provide new knowledge from experimental datasets. Our Community Phenotype Fitting method (CommPhitting), in contrast, balances resolution and flexibility by adjusting biologically meaningful variables and parameters while best recapitulating the provided dataset of the system. CommPhitting only requires two inputs – member metabolic models and dynamic growth data – and yields time-resolved predictions of metabolite concentrations, predicted abundances for each species and their expressed growth phenotypes, and linear growth rate constants for each expressed phenotype. These outputs offer broad value for a) understanding community biology and b) designing qualitative and quantitative metabolic hypotheses with mechanistic support. The contribution to hypothesis development specifically facilitates the judicious allocation of limited resources to experiments with computational evidence, thereby accelerating experimental and theoretical progress towards understanding microbiome dynamics. Dissonance between predictions and data can further be a source of actionable reflection that leads to experimental improvements, e.g. misaligned concentrations may elicit that the set of parameterized phenotypes are insufficient to explain the experimental data, which improves understanding of the community system.

We envision that CommPhitting can extend from the elucidation phase that we introduce herein to an exploration phase that is introduced in Step 4 of Figure 3. This exploration phase can test engineering hypotheses of community behaviors after member mutations and environmental perturbations, such as the presence of toxins/antibiotics. This exploration phase would fix the parameter and variable values that were derived from fitting experimental data in the elucidation phase and then would exchange the optimization objective to reflect ecological principles that guide the behavior of these systems (e.g. maximizing total community growth). The exploration phase may then come full-circle by looping into the elucidation phase as a design/build/test/learn cycle [69] where the mechanisms of an engineered community that was informed by the exploration phase are revealed.

The aforementioned unique attributes of CommPhitting enable it to resolve the abundances and linear kinetics of the growth phenotypes for each member of a microbial community more robustly than extant methods. This tool will be an invaluable resource for expanding basic knowledge of community dynamics and ultimately engineering these communities for applications in fields as diverse as bioproduction, medical therapies, and national security.

## Supporting information

Experimental metadata table

