## Supplementary material for "A microbial community growth model for dynamic phenotype predictions": Experimental metadata table

### Supporting Information: MSCommFitting

#### Contents

#### List of Figures

#### List of Tables

S1 The trials of batch experiments that were simulated by CommPhitting from our exemplar 2-member community. 2

| Experiment | Data | Trial simulated |
|---|---|---|
| Maltose | April 29, 2022 | B5 |
| Maltose+4HB | April 29, 2022 | F5 |
| Acetate | June 17, 2022 | E3 |

**Table S1.** The trials of batch experiments that were simulated by CommPhitting from our exemplar 2-member community.
